# Quantification of anticipation of excitement with three-axial model of emotion with EEG

**DOI:** 10.1101/659979

**Authors:** Maro G. Machizawa, Giuseppe Lisi, Noriaki Kanayama, Ryohei Mizuochi, Kai Makita, Takafumi Sasaoka, Shigeto Yamawaki

## Abstract

**Objectives:** Multiple facets of human emotions underlie diverse and sparse neural mechanisms. Amongst many models of emotions, the circumplex model of emotion is one of a significant theory. The use of the circumplex model allows us to model variable aspects of emotion; however, such momentary expression of one’s internal mental state still lacks to consider another, the third dimension of time. Here, we report an exploratory attempt to build a three-axial model of human emotion to model our sense of anticipatory excitement, “Waku-Waku (in Japanese),” when people are predictively coding upcoming emotional events.

**Approach:** Electroencephalography (EEG) was recorded from 28 young adult participants while they mentalized upcoming emotional pictures. Three auditory tones were used as indicative cues, predicting the likelihood of valence of an upcoming picture, either positive, negative, or unknown. While seeing an image, participants judged its emotional valence during the task, and subsequently rated their subjective experiences on valence, arousal, expectation, and Waku-Waku immediately after the experiment. The collected EEG data were then analyzed to identify contributory neural signatures for each of the three axes.

**Main Results:** A three axial model was built to quantify Waku-Waku. As was expected, this model revealed considerable contribution of the third dimension over the classical two-dimension model. Distinctive EEG components were identified. Furthermore, a novel brain-emotion interface is proposed and validated within the scope of limitations.

**Significance:** The proposed notion may shed new light on the theories of emotion and supports multiplex dimensions of emotion. With an introduction of the cognitive domain for a brain-computer-interface, we propose a novel brain-emotion-interface. Limitations and potential applications are discussed.

## 1. Introduction

Human emotions are complex and constructed of multiple facets of separable components. Amongst many models of emotion, a two-dimensional circumplex model comprised of valence and arousal axes originally proposed by Russell [1] is widely examined as a ubiquitous model across diversities of cultures [2,3]. Other theories, such as discrete categorical theory exist [4,5]; however, the majority of models agreeably assumes that our emotion is a product of a momentarily affective state. Recent developments in studies of emotion propose an update of the model of emotion. For instance, a ‘psychological constructionist approach’ conceptualizes emotion as an interplay of complex psychological operations, rather than the simplistically discrete emotional state [6]. Along the line, some studies conceptualize our emotion as an ‘affective working memory system’ that defines emotion as dynamic and active interactions between cognition and affect [7,8]. Based upon a concept that emotion innately interacts with our interoception of homeostatic state of the body[9,10], an alternative notion is also provided, such as emotion is driven by the result of internal inference of predictive coding [11,12]. As such, our affective awareness may be an interactive state between cognitive and emotional functions, and it may not be necessarily composed of a unitary function [see 13 for review].

These evidences suggest the importance of a novel, comprehensible psychological model of emotion as well as the corresponding neural decoder (also known as brain-computer-interface, BCI) to quantify our strikingly complex subjective experiences. Notably, the classical, oversimplified two-dimensional model may require an update as a model of our feelings. Recent trends in neurosciences and psychological sciences propose our brain as a predictive machine [11] supported by Bayesian theories on the human brain [14,15]. Such autonomous theory indicate that an emotional status might be explained with a sequence of instantaneous affective states to predict future states under uncertainty. In other words, one may feel being excited emotionally and at the same time, speculating on what to experience in the future cognitively. For instance, when we are excited by thinking about eating delicious food in a couple of hours, we are highly aroused and feeling happiness emotionally, and in the meantime, the brain may be automatically forming imagery of a particular food what one wants to have. Altogether, it is plausible that BCI approaches to quantify our subjective experiences may benefit from modeling our subjective feelings with the addition of the cognitive domain of anticipatory coding mechanism to the classical circumplex model of emotion. Here, based on the dimensional theory of emotion, we propose a 3-dimensional BCI model to quantify our mental experiences.

### 1.1 “Kansei”–a multiplex state of mood

Our motivation for this study originated from an idea to quantify a putatively complex state of mental representations, “Kansei.” In Asian languages, ‘*Kansei* (in Japanese, a direct translation would be ‘sensitivity’ or ‘sensibility’)’ is a widely accepted term that reflects one’s feeling that exogenously triggered by something and that often accompanied by mental images of a target [16]. *Kansei* expressions often reflect a mixture of affective and cognitive states, just as exemplified in recent views on human emotion [6]. For instance, being “Waku-Waku”, is one of the *onomatopoeias* states that is typically defined as an emotional state in which one is being excited emotionally while anticipating upcoming *pleasant* event(s) in the future cognitively. As the closest candidate of Russell’s circumplex model, an emotional state of ‘excited’ is placed between ‘alert’ and ‘elated’ with a moderate level of high valence (pleasant) and a moderate level of high arousal. Closest synonyms for “Waku-Waku” in English may be ‘anticipatory excitement’ or ‘a sense of exhilaration.’ As our eventual goal is to quantify such a putatively complex state, it was hypothesized the addition of an extra dimension of cognitive state might well explain the state of “Waku-Waku”. Therefore, we have hypothesized that a combination of *multi*-dimensional axes that incorporates both affective and cognitive dimensions, including prediction, could be a sensible model to reflect the state of *Kansei*. To note, in a broad sense, have *Kansei* not only a meaning for ones’ *state* but also reflects one’s *trait* or preferences based on one’s experiences. For the sake of clarity, we focus on the former *state* of one’s affect and cognition in this article. The other aspect of trait shall be treated elsewhere.

Here, we first aimed to build a psychological model for a *Kansei*, mainly focused on a state of “Waku-Waku.” Just like the expression “Waku-Waku,” *onomatopoeic* expressions are commonly used in the Japanese language to reflect an emotional state. Therefore, a neural-based quantification of *Kansei* would be of great advantage for many industrial applications. Based on a conventional two-dimensional model of affect [1], we hypothesized a three-dimensional model that was composed of ‘valence’ and ‘arousal’ axes as well as the third dimension of prediction, ‘expectation.’ Please note, it was defined that ‘expectation’ only has cognitive aspect of anticipation putatively forming explicit imagery of a future under a situation of uncertainty. It may provoke a degree of emotional valence; however, such potential overlap was treated mathematically (see Methods). Whereas by definition, ‘Waku-Waku (anticipatory excitement)’ holds emotional valence and arousal when expecting a pleasant event in the future as described above; therefore, it was expected to be loaded by both affective and cognitive states of mind.

### 1.2 Brain-computer interface

Recent developments in neurosciences allow us to build an interface to monitor our neural status in real-time by presenting its status on a screen or a device. Today, the demand for techniques such as neurofeedback or BCI is increasing in clinical settings and even in industrial applications [17-19]. Several variations of neurofeedback methods exist; some apply to visualize one’s neural state by using functional Magnetic Resonance Imaging (fMRI) for training and therapeutic purposes [20,21]. BCI with electroencephalography (EEG) has also been widely employed to detect a locus of attention with the P300 component [18,19] or as a means to assess conscious level with *α*-waves [22-24], and so forth. These classical models typically acquire data from one or a few electrode channels and focus only on a specific frequency range of interest. With improved computational resources, BCIs focusing on limb movement acquire real-time feedback of independent component (or common spatial pattern) recorded from the whole scalp electrodes [25-27], rather than data from solely on one or few channels that was common in the past. Nowadays, one can effortlessly expect a further complex BCI could be achieved, such as building a neurofeedback system incorporating multiple neural indices such as the multi-dimensional model proposed here.

### 1.3 Brain-emotion interface

In this study, we first modeled “Waku-Waku” with three dimensions: namely valence, arousal, and expectation provided the psychological model for “Waku-Waku” as an intermixed state of higher-order affective and cognitive functions [7]. Having confirmed the significance of all axes in the psychological model, secondly, we derived electrophysiological markers using EEG corresponding to each axis. At last, based on the outcomes, we propose a linear equation model of “brain-emotion-interface (BEI)” that may be able to quantify “Waku-Waku” by incorporating the three-dimensional psychological model with corresponding neural markers for each of the three axes in real-time. Below, we show the resultant psychological model, EEG markers for each axis, and propose a prototype 3-D model for the quantification of “Waku-Waku.” Potential applications of the BEI as a tool of *Kansei*-engineering and limitations in this study are discussed.

## 2. Methods

As the first step, we focused on building a psychological model for the “Waku-Waku.” We performed a picture rating experiment in which participants were asked to imagine what kind of new picture would be displayed depending on a received valence-predicting cue. Immediately after the main task, participants completed a subjective rating task to evaluate their feelings during the task as well as emotional responses for each picture. Details of the experiment and analyses are as it follows.

Given the hypothesis, an original experimental plan was to elucidate brain functions with fMRI as well as EEG, thereby capturing multi-modal scopes of *Kansei*. The same participants visited the lab three times, twice for fMRI sessions, and once for an EEG session. The part of fMRI outcome has been reported elsewhere [see 28]. The same task was repeated three times, once with an EEG recording and twice with fMRI recordings. We report the fMRI session below because it was necessary to include the subjective rating data obtained from all three visits of 28 participants to derive a satisfactory linear model.

### 2.1 Participants

Thirty-six healthy young adults (19 females) aged between 19–27 years old were recruited locally. Due to technical errors or early termination of all three visits to the lab, some participants were rejected. As a result, data from 28 participants (16 females; age mean ± *SD*: 22.17 ± 1.79) are reported in this report. All reported no history of neurological or psychological disorders. All participants had normal hearing abilities with either normal or corrected-to-be normal vision. All participants gave their informed consent approved by a local research ethical committee located at the Hiroshima University.

### 2.3 Behavioral procedures

Participants performed a picture rating task in which they were requested to mentalize upcoming novel picture appearing on a computer monitor. One of three variants of auditory cues preceded a picture onset (see Figure 1). These 3 cueing conditions were: 1) a low-tone predicting a positive picture (‘Predictive Pleasant’), 2) a high-tone predicting a negative picture (‘Predictable Unpleasant’), and 3) an intermediate tone indicated a probability of seeing a pleasant and unpleasant picture was 50-50 (‘Unpredictable’). An auditory tone was played for 250msec, followed by a blank delay period for 4,000 msec, including the initial 250msec tone. The conditional assignments for the high and low tones were counterbalanced across participants. This assignment was fixed throughout all of the three visits. While arousal of the pictures varied picture by picture, these auditory cues were unrelated to the arousal of the upcoming image. To note, we categorized each picture for the purpose of counterbalancing picture sets across different sessions; we referred to subjective ratings of valence and arousal reported in the original article [IAPS; 29] (Supplementary Figure 1 and Supplementary Table2 for detailed ratings of valence and arousal. Appendix lists all picture codes used in this study). Participants also rated based on their subjective feelings after the experiment (see below for details).

**Figure 1.**
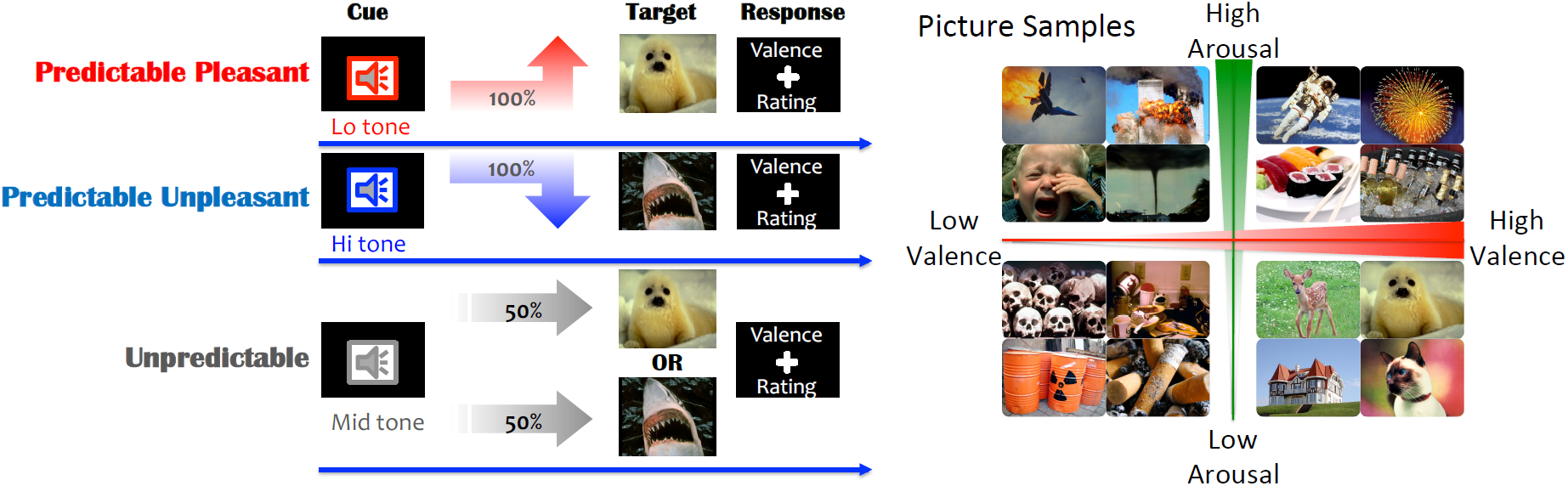
Experimental schematics of the task (left panel) and sample IAPS pictures selected in this study (right panel). Participants heard one of three conditioned auditory tones, each of which predicted upcoming picture to be a positive (‘Predictable pleasant’), negative (‘Predictable Unpleasant’), or either of those at 50% probability (‘Unpredictable’). Pitch of tones indicated which condition that trial would be. A low tone, an intermediate tone, or a high tone corresponded with positive, unpredictable, or negative conditions (assignment for the high and low tones were counterbalanced across participants). After a 4,000 msec inter-stimulus-interval (ISI), including 250msec of the tone, a picture was displayed for 4,000 msec followed by a valence rating task after an image offset. During the ISI, participants were requested to imagine what type/kind of picture might be displayed. At the rating period, participants were required to rate emotional valence of perceived picture at a 4 Likert-scale, ranged the most negative, more-or-less negative, more-or-less positive, to the most positive. The right panel shows sample pictures displayed in this study, mapped onto quadrants of valence and arousal axial model.

During the 4,000ms blank period, participants were requested to imagine what kind of image would be displayed. After the delay, an emotional-triggering picture was displayed at the center of the screen for 4,000ms (for the details of the selected pictures, see Stimuli section below). Followed by the picture display, a red fixation cross appeared at the center of the screen for 1,000ms. During this response period, participants were requested to rate their subjective feeling of valence, how they are moved by seeing that picture (on a 4 Likert-scale: ‘strongly pleasant’, ‘pleasant’, ‘unpleasant’ and ‘strongly unpleasant’) by pressing a corresponding button with a thumb, index finger, middle finger, or a ring finger on a keyboard (for EEG) or a response button (in fMRI). Before the main task, participants underwent a brief practice session for all three conditions with a picture set independent of the main task.

### 2.3 Stimuli

As a predictory cue, one of three different auditory tones (500, 1,000, or 1,500Hz) was played for 250 msec via a headphone worn comfortably. Before the experiment, during the practice session, volumes of tones were checked with participants and adjusted to their comfortable hearing-level. Either 500 Hz (a low tone) or 1,500 Hz (a high tone) predicted either pleasant or unpleasant, and 1,000 Hz (an intermediate tone) was used as an unpredictable cue. The assignment of high and low tones was counterbalanced across participants. The intermediate tone was fixed as the unpredictable condition across all participants.

As novel images, pictures were carefully selected from 1182 the International Affective Picture System [IAPS; 29] with the following criteria. Pictures that might cause an excessive negative affect, such as corpses or alike, or that may interfere against our local ethics were discarded. In addition, pictures consisting of multiple objects where people may not necessarily focus on one aspect of that picture, pictures with intermediate valence that may not evoke adequate intensity of valence either positive or negative (such as a plain scenery or object; with valence ratings between 4–6, see Supplementary Figure 1), pictures containing texts or symbols or items that may have cultural discrepancies (i.e., guns) were disregarded for our picture sets. The resulting 320 pictures were divided into 2 sets of 160 pictures (one for two-times of the test for MRI sessions with 80 trials each and the other 160 pictures for an EEG session) randomly by controlling for average ratings (as reported in the IAPS dataset) of valence and arousal. Each set of 160 pictures were counterbalanced across participants. Of 160 pictures, half of them were pleasant, and the other half was unpleasant. Each picture appeared at least once for a predictory cue (predicting pleasant or unpleasant), and a half of the 80 pictures each (pleasant and unpleasant) were used twice as a condition of an unpredictive cue (‘intermediate tone’). Of selected pictures, spatial frequency and brightness were also controlled for the picture sets so that brightness and spatial frequency (high or low split at 140 Hz) of visual features were not significantly different among different sets. Besides, the contents of each picture have been visually determined (human, animal, scenery, or others), and the categorical information was also equally distributed into each set. See Supplementary Figure 1 for the details of valence and arousal ratings used in this study. All IAPS pictures were novel to the participants.

A Dell 24-inch LCD monitor was used to display pictures at a 1920 × 1080 pixels resolution. A chin-rest placed 56cm away from the monitor was used to stabilize monitor to the eyes distance across participants. Participants were asked to rest their chin during the task. The size of pictures varied; some oriented in landscape, and others were in portrait; however, the original pictures were displayed to fit the monitor. All auditory and visual stimuli were delivered by the Presentation software version 17.2 (NeuroBehavioral Systems, San Francisco, USA). As noted above, there were fMRI and EEG sessions. All behavioral tasks remained the same, except that the number of trials differed for the fMRI sessions; a total of 120 trials for an fMRI session, instead of 240 for EEG, were performed. On each visit, different sets of pictures were used to maintain the novelty of pictures. For a case of the EEG session, there were 80 trials for each cueing condition and a total of 240 trials for an experiment. Regardless of cueing type, participants judged 120 pleasant and 120 unpleasant pictures.

### 2.4 Subjective rating procedures

At the completion of the main task, participants reported their subjective feelings by rating on a 0–100 visual analog scale for all conditions. Each question was displayed at the center of the screen, and a VAS was displayed underneath. Participants answered each question on a computer monitor by moving a pointer to the left (0) or the right (100) on the scale by pressing left or right arrow keys on a keyboard. Participants were instructed: “The following questions are about the evaluation of your feeling during the experiment. Once a question appears on the screen, please answer the degree of feeling when you heard each tone during the experiment on a scale between 0–100”. For each condition, participants rated the degree of ‘valence’ (unpleasant to pleasant; ‘Kai’ in Japanese), ‘arousal’ (low to high arousal; ‘Kassei’ in Japanese), and ‘expectation’ (low to high expectation; ‘Kitai-kan’ in Japanese), as well as “Waku-Waku” (in Japanese, exhalation or excitement). For example, the sentence below appeared in the middle of the screen for the ‘predictive pleasant’ condition: “Please rate the degree of “pleasure (wellness of the feeling)” in which you heard a low-tone.” As for arousal, “arousal (degree of liveness),” and as for expectation, just “expectation.”

Here we selected the term ‘expectation,’ rather than a word, ‘prediction’ or ‘time.’ It was because 1) we assumed such predictive sense would be hard to report consciously and accurately, 2) ‘time (how soon they would feel/expect)’ was inappropriate to ask as the timing of upcoming event of seeing a picture was fixed to 4 secs. While the definition of ‘Waku-Waku’ might be depended on how one would conceptualize it. For the sense of anticipatory excitement, the term ‘expectation (Katai-Kan in Japanese)’ was supposed to reflect anticipation. However, to note, one might not be able to adequately separate and ignore their own emotions covertly attached to the expectation when rating the expectation. As their feeling of expectation might unconsciously induce emotional valence. We discuss the interpretation with the derived correlation in Results Section 3.1 and Discussion; however, our choice of statistical procedure of the mixed model would account for such correlated inputs (see below for details).

As described above, each participant rated a total of three times for each visit of the experiment (once for EEG, twice for fMRI experiments). All rating results have been accrued for the analysis. To note, building a model with 28 participants’ data only from one EEG experiment (one data point per condition) did not suffice our statistical criteria to construct a psychological model. We decided to include the entire three sessions’ data to assure all three axes achieved significance level.

### 2.5 Statistical procedures

A *mixed* linear model was computed based on these subjective ratings to model the anticipation of excitement, using SPSS version 22. Compared to the general linear model (GLM) that assumes independence across data, the mixed linear model has an advantage in handling putatively correlated and unbalanced data. Notably, some degrees of correlations and random-effect among subjective ratings across axes were assumed; a typical GLM approach might not be suitable for this case. The resultant psychological model, therefore, takes putatively correlated effects into account. As we aimed to model Waku-Waku feeling with variants, the anticipation of excitement (“Waku-Waku”) was a dependent variable, and ‘valence,’ ‘arousal,’ and ‘expectation’ were independent variables. Assignment of counterbalanced tones, used picture sets, as well as examined domain of measures (two MRI measures and one EEG measure) were included as covariates of *no* interest.

As described above, the inclusion of at least valence and arousal of the original circumplex model was expected to be fundamental to the model. To validate observed coefficients, the likelihood ratio test (χ2 statistic) was evaluated on each axis. All axes met the significant level (*p* < .05) reported formula in section 3.1. As it turned out, the arousal axis did not meet the criteria when including only one of three sessions. That was a reason to include behavioral results from MRI sessions that could have been an independent study.

### 2.6 EEG procedures

#### 2.6.1 Recording procedures

During the task mentioned above, participants’ EEGs have been recorded with 64ch BioSemi Active Two system at a sample rate of 2,000 Hz. Channels were placed according to the International 10-20 system montage. In addition, a vertical and a horizontal electrooculograms were collected as in convention (approximately 3–4cm below and above the center of the left eyeball for vertical, and approximately 1 cm horizontally to the side of external canthi on each eye). An online reference channel was placed on the tip of the nose.

#### 2.6.2 Analytical Procedures

Recorded EEG data were analyzed offline with the EEGLAB toolbox [30] running on the Matlab 2015a (Mathworks, Inc), and it was partly combined with custom-made functions. Continuous data were first removed its DC-offset, low-pass filtered with two-way least-squares FIR filter at 40 Hz, resampled to 512Hz, epoched from 500 msec before cue onset to 8,200 msec after the cue onset (4,200 msec after image onset). The epoched data were then average re-referenced, and each channel was normalized to the baseline period (the 500 msec before the cue-onset). Any trials with excessive artifacts on channels were rejected by the automatic artifact rejection model implemented in the EEGLAB with an absolute amplitude threshold with more than 100µV, probability over 5 standard deviations. Each iteration of the artifact detection was performed with a maximally 5% of total trials to be rejected per iteration. In addition to the basic artifact rejections, we corrected artefacts derived from eye movements using conventional recursive least squares regression (CRLS) implemented in the EEGLAB [31] by referring to the vertical and horizontal EOG reference channels with 3rd order adaptive filter with a forgetting factor (lambda, ‘λ’) of .9999 and .01 sigma (‘σ’).

The resulting corrected data received the first run of independent component decomposition (also known as, ‘ICA’) [30] to capture any artifactual activities that still survived our initial rejection criteria. To be specific, logistic infomax ICA algorithm [32]–– an unsupervised learning algorithm and a higher-order generalization of well-known principal component analysis to separate statistically and temporary independent components in the signals–– was used. It was followed by another run of automatic artifact rejection now on the independent components (ICs) to remove artifactual components with the same criteria used for the channel-based rejection, as mentioned above. After the IC-based rejections, the artifact-free data underwent the second, and the last IC decomposition to extract independent neural activities this time, resulting in 64 putatively clean ICs per participant. Finally, fine-grained dipole analysis was performed using the dipfit function in the EEGLAB for each IC assuming one dipole in the brain. Supplementary Figure 2 depicts the estimated dipole positions for each IC.

#### 2.6.3 Rejection criteria

Any ICs with residual variances more than 50% (equivalent to proportion of outliers at *p* < .005, one-tail; z-score > 2.58), estimated dipole positions outside of the brain, or any ICs with an inverse weight only on one of EEG channel with more than 5 standard deviations among the rest of channels that is likely to be an artifact of the channel were rejected. This process retained an average of 33.89 ICs (ranged between 26–43 ICs) per participant. A total of 949 ICs was retained for subsequent analyses.

#### 2.6.4 Clustering independent components

In order to quantitatively determine the number of IC clusters to be extracted, we employed a gaussian mixture model (GMM) to cluster ICs based on their scalp topography, and we iterated GMM across a range of potential number of clusters (1 up to 60), and the number of clusters to extract was determined by Bayesian Information Criteria (BIC) due to its consistency over Akaike Information Criteria [33,34]. Because of the nature of ICA, the polarity of the IC scalp map is arbitrary. Therefore, polarity of all retained IC was aligned by inverting polarity of each IC weights, if necessary, such that all components correlate positively to each other *prior to* the computation of the gaussian mixture models. All the aligned data were then Z-score normalized across channels per IC before the GMM analysis.

We iteratively clustered the 949 ICs with their inverse weights of the 64 channels by a gaussian mixture model that maximizes likelihood using the iterative expectation-maximization (EM) algorithm with the following rules. Covariance type was restricted to be diagonal; shared covariance was allowed, and with an addition of regularization value of 0.05. A maximum number of allowed EM iterations within each fit was set to be 1,000. We repeated the procedures for 1–60 clusters (we did not perform more than 60 as the decision could have been drawn straightforwardly from this number). The best GMM was determined by their BIC values. Finally, centroids of inverse weights for each cluster were computed, and then each IC was clustered based on the selected model for subsequent statistical analyses.

#### 2.6.5 Spectrogram computation

The pre-processed data was re-epoched from –500 to 4,200 msec around the *image*-onset for valence (seeing positive v.s. negative pictures) and arousal (seeing high v.s. low arousal pictures) and baseline was corrected between –500 and 0 msec. Likewise, data was re-epoched from –500 to 4,200 msec around the *cue*-onset for expectation (expecting positive picture v.s. unpredictable), and baseline was corrected between –500 and 0 msec. Please note, both epoched data shared the same ICs as this epoch separation was done *after* the final ICA so that we can equally compare the results before and after the image onset.

For all retained ICs, spectrogram was computed between 0–4,000 msec from the onset of *image* for ‘valence’ and ‘arousal,’ and 0–4,000 msec after the onset of *cue* for ‘expectation.’ For the sake of practicality with its speed of computation in real-time, we applied Fast Fourier Transformation (FFT) on each segmented data with a hamming window and zero-padded ratio to assess the frequency power density in each band. The resulting spectral power was then averaged for each frequency range of *θ* (4–8 Hz), *α* (8–12 Hz), and *β* (12–20 Hz). One of the purposes of this study was to elucidate neural correlates that would be efficiently applicable for academia-industry collaboration, under an assumption that a wearable and dry-electrode EEG headset might be used in various environmental situations. Compared to the active and wet EEG electrodes with a low electrode impedance what we have acquired in this study, dry electrode headsets are known to be susceptible for low signal to noise ratio [35], particularly at slow and fast frequency bands including event-related potentials. Moreover, a higher electrode impedance also is a cause for the sweat that strikingly diminishes data quality depending on the recording environment, especially in a hot and humid environment [36]. Therefore, we have focused only on these frequency bands that are likely to be reliable frequency bands with potential easy-to-use applications in mind.

Finally, we examined whether a spectral power of each IC can dissociate each type of valence, arousal, and expectation processes. Because spectral powers for each IC did not distribute for most cases, Wilcoxon signed-rank test was applied, and its alpha-level was corrected by false discovery rate (FDR) method, controlling for multiple comparisons across frequencies.

#### 2.6.6 Evaluation of proposed BEI model

Among the three cueing conditions, our hypothetical choice for the third axis was set to only one of paired comparisons as the expectation axis. As one may speculate, an alternative contrast could be proposed, and it requires validation. Additional analyses were performed on ‘predictive pleasant v.s. predictive unpleasant’ and ‘both predictory trials (predicting pleasant and predicting unpleasant) v.s. unpredictable’. The former contrast assessed feasibility against the valence axis because of sharing a similar contrast before and after the image onset. This contrast might have shared some neural properties on its emotional values even before seeing the image. The latter contrast was examined as a counterpart for the expectation axis, provided we can assume expectancy regardless of emotional values as the third axis. The choice of contrast, ‘predictive pleasant v.s. unpredictable’ as the expectation axis, was based on the way we asked participants to rate ‘expectation.’ The term might reflect not only predictability of an event (‘how likely to expect’) but also a positively biased degree of the mental imagery associated with the future (‘how much to expect’). Because the probability of seeing a pleasant/unpleasant picture was radically fixed at 100% for the predictive cues, collecting subjective feedback for the likelihood (or predictability) might be redundant in this experimental design. Therefore, it was assumed that any neural features corresponding just “prediction” might fail to reflect ‘expectation’ that might be biased to positive anticipation.

As it turned out, we found corresponding EEG features for each axis of interest. As a mean to validate the BEI model based on the EEG spectral powers, the proposed model was evaluated by estimating scores of each axis and then applied to the formula (4) (see Section 3.3 below). At first, spectral powers of selected frequency band of selected ICs in all conditions were converted into normalized scores (0–100) for each IC using the normal cumulative distribution function to match spectral powers (in a unit of decibels) to the VAS-scaled scores of Waku-Waku. For each axis, the obtained mean scores were then normalized across the 3 anticipatory conditions (predicting pleasant, predicting unpleasant, and unpredictive conditions) in which participants rated Waku-Waku. Please note, neural markers for valence and arousal axes were established based on EEG data *after* the image onset, but we have intuitively applied these neural markers during the anticipatory period so that quantification of Waku-Waku could be achieved based on the same EEG data as the expectation axis. At last, Waku-Waku was estimated by submitting converted scores (0–100) into the formula (4). Similarly, as a supplement to what the proposed BEI model could estimate, the transition of Waku-Waku was also quantified *after* the image onset period. As a comparison counterpart of ‘expectation’ axis, each conditional value was estimated by the ‘prediction’ axis, as mentioned above.

## 3. Results

### 3.1 Psychological model of “Waku-Waku”

See Figure 2 for a summary of subjective ratings for each cueing condition (‘expecting pleasant,’ ‘unpredicting,’ and ‘expecting unpleasant’). Based on these ratings for valence, arousal, expectation, and Waku-Waku, a mixed linear model was tested with valence and arousal as independent variables for a conventional 2-dimensional model and with valence, arousal, and expectation as independent variables for a 3-dimensional model. For both models, Waku-Waku was a dependent variable.

**Figure 2.**
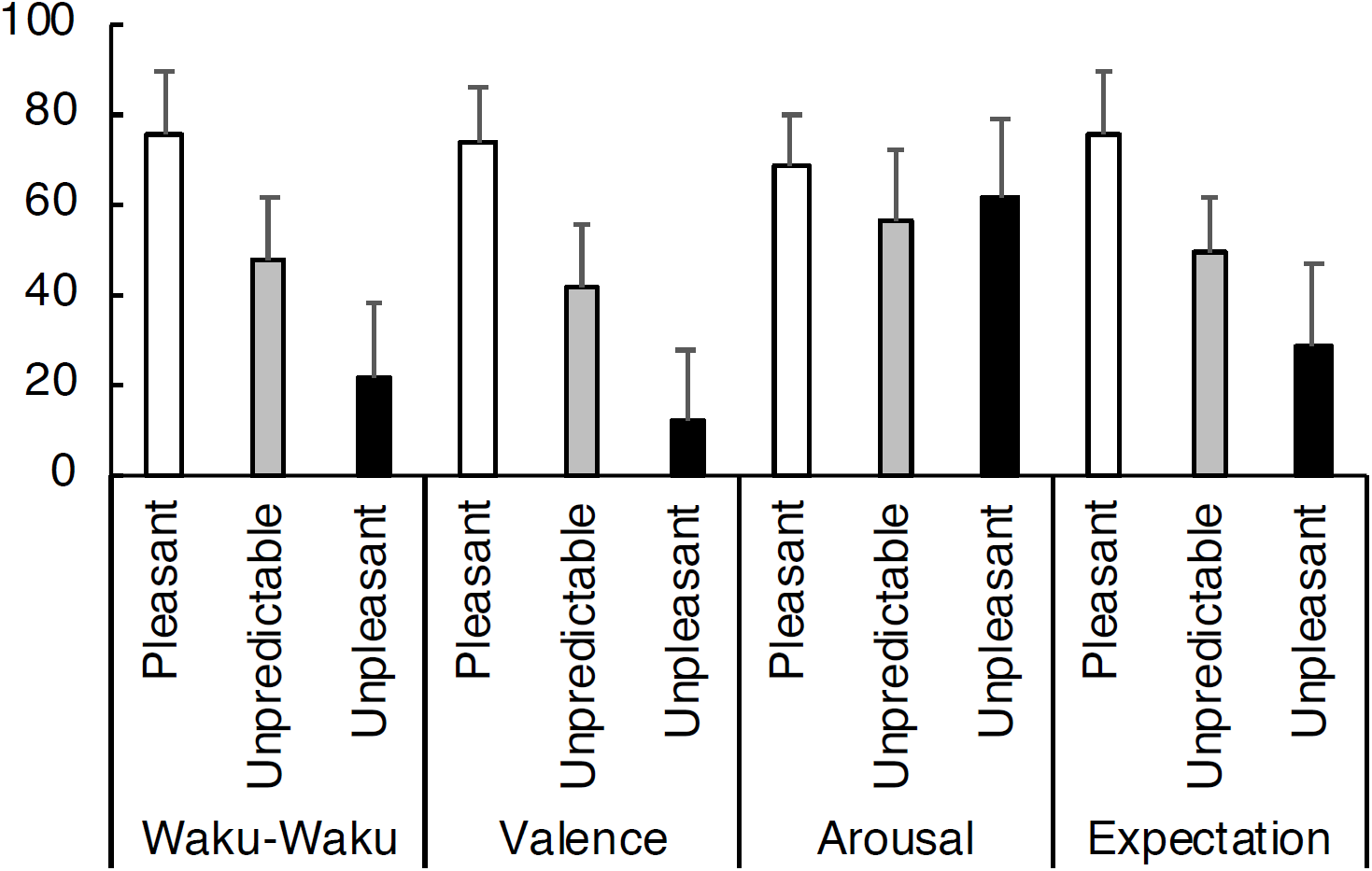
Grand-averaged subjective ratings of anticipation of excitement (“Waku-Waku” in Japanese), valence, arousal, and expectation on a 0–100 scale for each cued condition: namely, predictive pleasant (‘Pleasant’), unpredictive of either pleasant or unpleasant (‘Unpredictable’), and predictive unpleasant (‘Unpleasant’). These grand-averages were comprised of all three sessions per participant. Error bars represent 1 *SD*. As was expected, Waku-Waku was the highest for the predictively pleasant condition, followed by unpredictable and predictive unpleasant conditions. Ratings for valence were more or less similar to the anticipation but not necessarily the same. Arousal was rated equally across conditions. Expectation (see the main text for definitional differences between Waku-Waku and expectation) was similar to that of anticipation of excitement. A mixed linear model was performed on these subjective rating scores to model the anticipation of excitement. The resulting formula is reported in the main text.

With the 2-axial model, “Waku-Waku (‘W’)” was modeled as follows (adjusted *R*^2^ = .90):

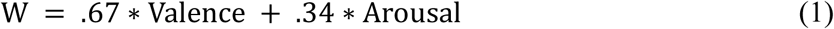

The linear model for the 3-axial model was as follows (adjusted *R*^2^ = .93).

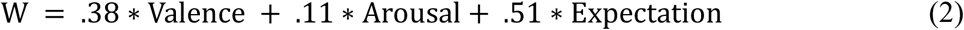

As was expected, the 3-dimensional model topped its fitting accuracy by 3 percent of the variance. Notably, the added third axis of expectation was significant and highly loaded. When including only one of experimental sessions, a coefficient for the arousal axis did not meet our criteria (at *p* < .05), but the other two valence and expectation axes did (*p* < .05). For completeness, here are the fitted formula for each session: [MRI1: W = .53*V + .04*A + .45*E; MRI2: W = .30*V + .12*A + .52*E; EEG: W = .29*V + .16*A + .61*E], where W, V, A, and E correspond to Waku-Waku, valence, arousal, and expectation, respectively. Akaike Information Criteria (AIC) values, an index to assess goodness-of-fit of each model, were compared for each session (MRI1, MRI2, and EEG) with the three axial models were: 670, 686, and 669. This result implies that the formula for the EEG session might have been the best compared to MRI sessions. However, the value differences of the three were so small and negligible, and these AIC results validate comparability of the scores on all three sessions.

In addition, direct pairwise correlations were examined between Waku-Waku and each of three axes (*r*2-values and p-values in parentheses) with valence, arousal, and expectation were .59 (*p* < .001), .04 (*p* < .005), and .66 (*p* < .001), respectively. These direct correlations roughly reflect weight balance of the coefficients derived in the formula (2). Correlations among the three axes were also tested: valence and arousal, valence and expectation, and arousal and expectation were: .00 (.37), .55 (< .001), and .05 (< .001), respectively. Please remind, the mixed model should have taken care of this correlated variables as well as the other unbalances or random-effect of samples. See Supplementary Figure 2 for scatter plots across axes.

### 3.2 Spectral EEG markers

As a result of the GMM on the inverse weights of 949 ICs, BIC criteria comparison between 1– 60 clusters suggested to aggregate 15 clusters. Figure 3 shows BIC values and clustered IC maps (Supplementary Figure 3 also depicts centroid coordinates of dipole location for each cluster).

**Figure 3.**
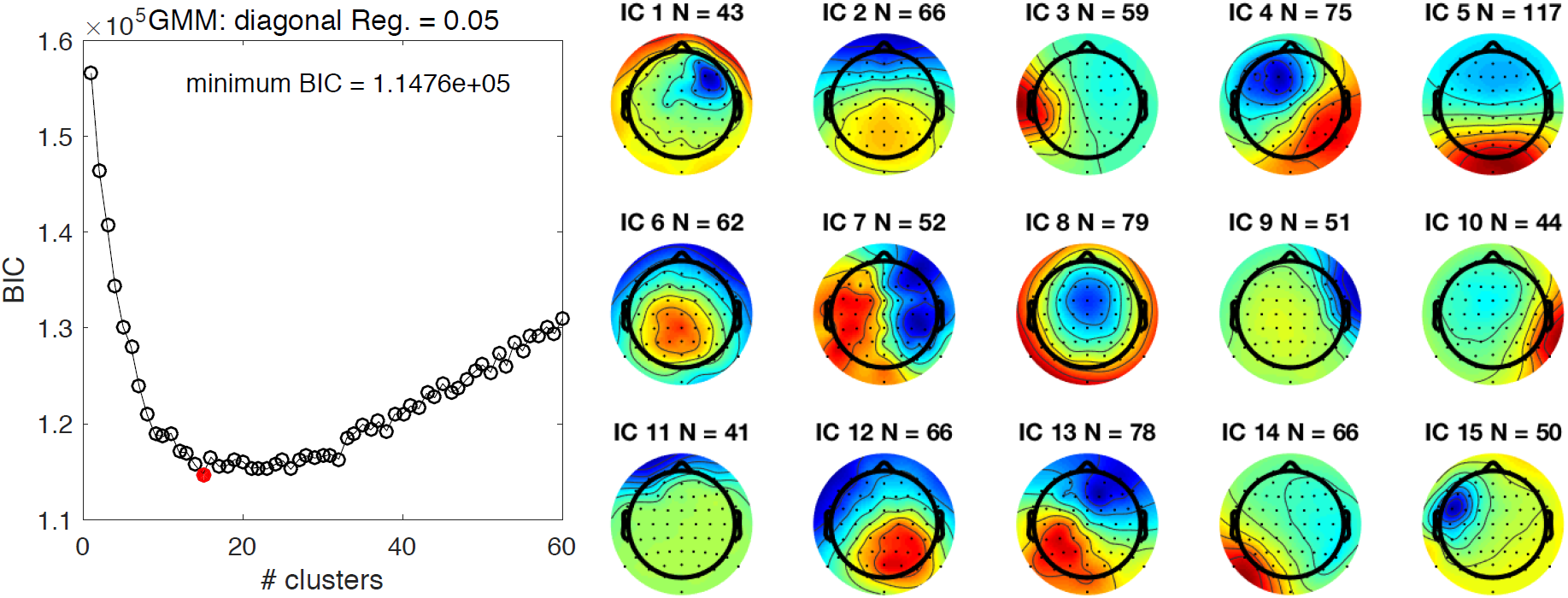
Bayesian information criteria (BIC) values of the gaussian mixture model (GMM) for each number of clusters (left panel) and 15 scalp topography of independent component (IC) clusters determined by the GMM (right panel). The number of clusters to extract was determined on BIC values. As it turned out, a model with 15 clusters (a red dot) was determined from 28 participants with 64 ICs per participant. “N” corresponds to the number of ICs grouped in the cluster. For more details, see Supplementary Materials. Note, cluster number is not important here.

Spectral power has been examined for each IC cluster (see Figure 4 for a summary of significant and marginally significant ICs; Supplementary Table 1 contains the statistical results of all IC clusters for completeness). For valence, arousal, and expectation axes, 5, 2, and 2 IC clusters emerged as significant, respectively. Emotional valence was represented by several EEG features, such as *θ*-band of IC clusters 1, 5, and 7, *α*-band of IC clusters 5 and 6, and *β*-band of IC clusters 7 and 14. Of these, IC cluster 5 was shared by all sampled participants. Arousal instead was quantified by two IC clusters: α band of IC cluster 7 and *β* band of IC cluster 11. Of these, the *α*-band of IC cluster 7 was more reliable (shared by a higher number of participants), shared by 86% of participants. As for expectation, *the θ*-band of IC cluster 10 emerged to be significant, and it was slightly more reliable compared to another IC cluster 15.

**Figure 4.**
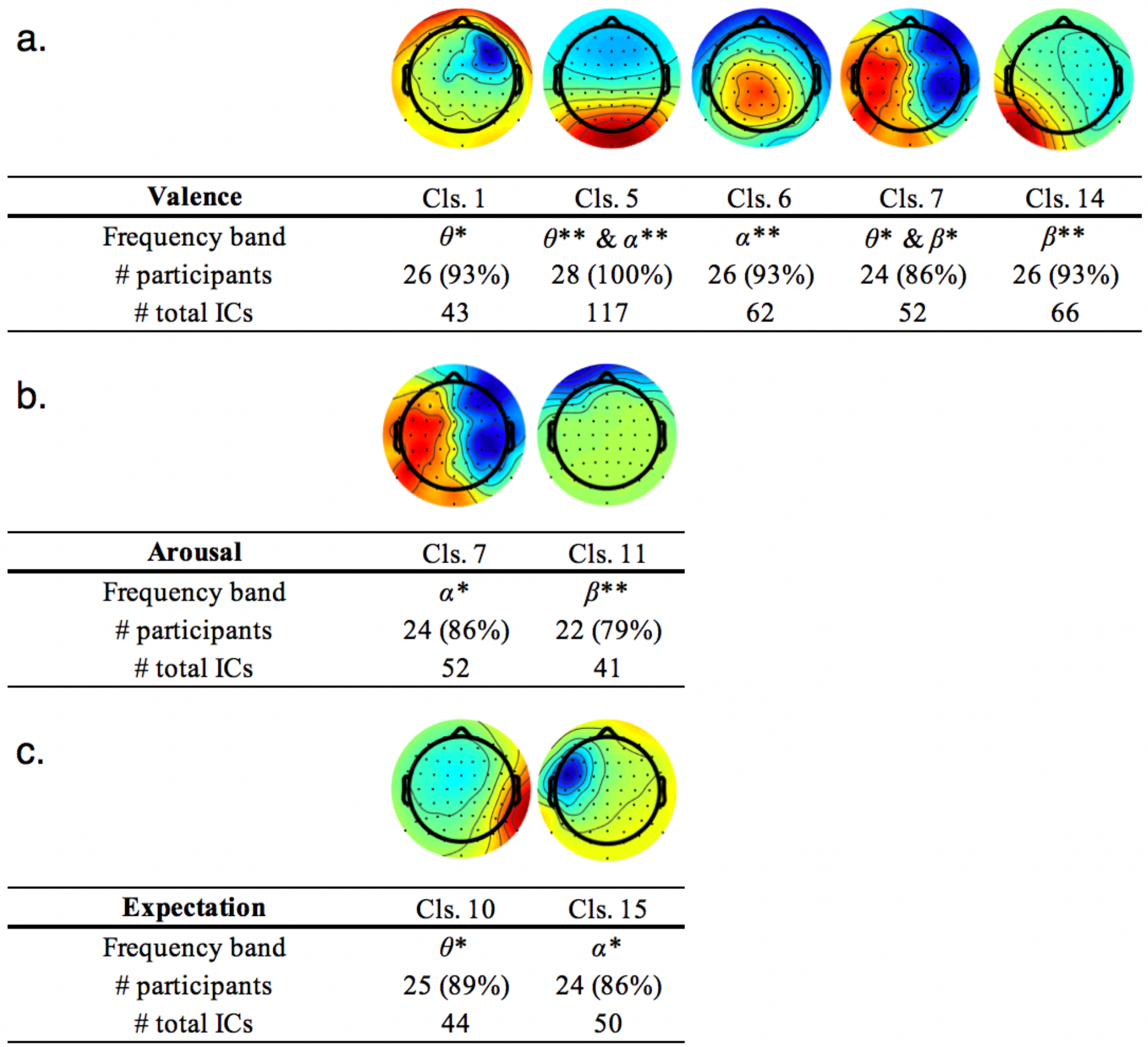
Scalp topographies of independent component clusters and their frequency ranges found to be significant for a) *valence* axis (seeing positive picture v.s. seeing negative picture); b) *arousal* axis (seeing high arousal picture v.s. seeing low arousal picture); and c) *expectation* (anticipating positive picture v.s. unpredictable, 50–50 chance to see positive picture). The number of participants who held an IC grouped into each cluster (‘Cls.’), and its percentage of participants relative to the sample size (*n* = 28) are listed as well as a total number of ICs belong to each cluster. Color intensities of either red or blue indicate a strong weight on the region. For our purposes, their red/blue color representation (either positive or negative) is not relevant as our focus of the analysis was solely on spectral power without consideration of the direction of the current. The color spectrum of red to blue may be flipped to represent the same IC. **Statistically significant at *p* < .05 FDR corrected; *Statistically significant at *p* < .05 uncorrected. For the full details of the statistics for all clusters, please refer to Supplementary Table.

Supplementary Table 4 shows the results of all ICs, including marginally significant ICs, as well as additional supplementary comparisons focusing on ‘predictive pleasant v.s. predictive unpleasant’ and ‘both predictive conditions v.s. unpredictive’. Some IC clusters were shared by all participants (100% of them), but some were not (i.e., cluster 11 for valence with 79%). Across all comparisons, some IC clusters are shared on different axes, such as cluster 7 for valence and arousal at different frequency bands. No single IC cluster (and its same frequency range) survived the correction across all the three axes.

### 3.3 3-D linear model of BEI

Given the three-dimensional psychological model (2) and corresponding neural correlates selected for each axis, we first propose a ‘conceptual’ BEI model to estimate Waku-Waku (‘We’) below. Assuming that the formula (2) is valid and that corresponding EEG features could estimate each axial value (ranging between 0-100), the psychological axes in the formula (2) consisted of subjective ratings may be replaced with corresponding EEG features.

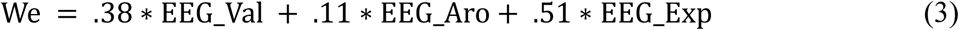

where EEG_Val, EEG_Aro, EEG_Exp correspond to normalized (0–100) spectral power of an IC for valence, arousal, and expectation axes, respectively. Here, a conversion of EEG spectral power in its unit (Db) to a normalized unit (0–100) is necessary because the unit acquired for the psychological model ranged between 0–100 while spectral density of EEG signals does not range between 0–100.

Given the statistical results, we selected the most robust and reliable IC candidates that survived our criteria because multiple EEG features were identified. The selection criteria were a combination of the following: the highest Z-score and the highest proportion of participants who held the selected IC cluster. The final selected formula for the estimated Waku-Waku (We) is expressed as below:

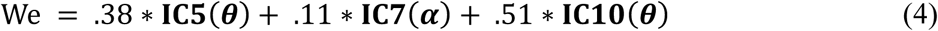

where IC5, IC7, and IC10 correspond to IC cluster reported in Figure 3, Figure 4 and Supplementary Table 1; *θ* and *α* in parentheses correspond to the frequency range of interest for each IC. Please note, as described in the above section, each EEG feature assumes the values are normalized (0–100) within its power distribution. An example workflow using the formula (4) is depicted in Figure 5.

**Figure 5.**
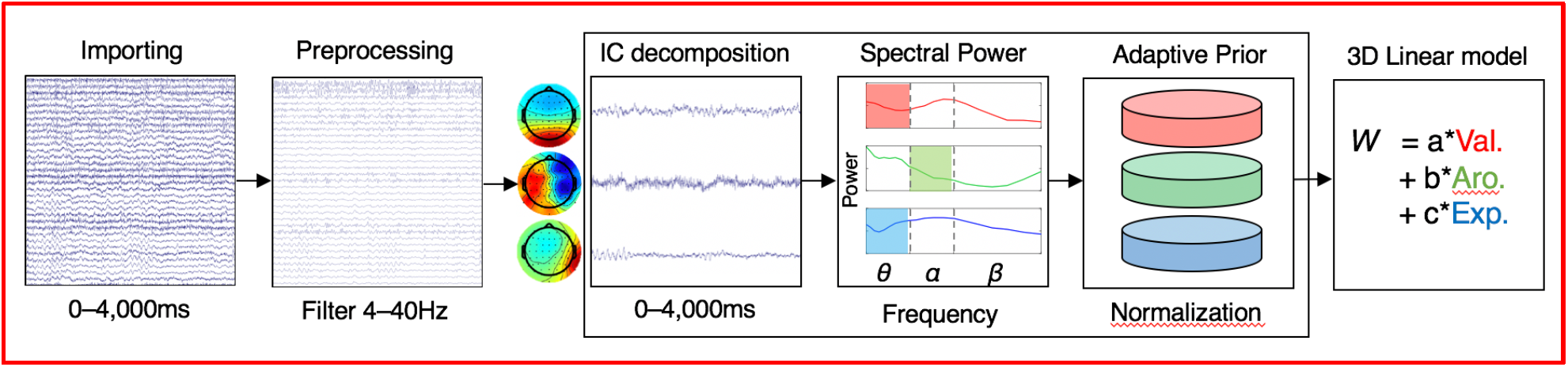
A conceptual workflow of quantification of “Waku-Waku (anticipatory excitement, ‘W’)”.

In Figure 5, we propose an example BEI workflow. First, raw EEG data of a chunk of 4,000ms would be imported. The imported data would be first preprocessed (including filtering 4–20 Hz and data mining) for subsequent quantification for valence, arousal, and expectation axes. IC weights of interest (IC5, IC7, and IC10 as in Figures 3 and 4, respectively) would be extracted on sphered data (this is a necessary step to compute IC), spectral power density for each target frequency (*θ* of IC5 for valence, *α* of IC7 for arousal, and *θ* of IC10 for expectation, as were determined in Section 3.2) would be obtained, and the obtained spectral power is then scaled in a normalized unit (0–100) based on the prior distribution of the spectral power of each IC candidate. The final step simply combines the coefficients for each axis (as in formula (4)) with those normalized EEG features. The prior distribution of each EEG feature may be assessed on the entire collected data for offline analysis. Alternatively, for a case of real-time workflow, the prior distribution may be constructed by collecting a certain duration of data as a calibration stage prior to run this. With the rescaled EEG features, the final values for valence (‘Val.’), arousal (‘Aro.’) and expectation (‘Exp.’) are replaced by those values obtained from EEG as in formula (4).

### 3.4 Validation of the neural markers and the 3-D linear model

While we proposed a hypothesis-driven BEI model to conceptualize and quantify Waku-Waku with three theoretically separate axes and corresponding neural markers, it would be still necessary to validate our proposed model with the existing data samples even if a robust interpretation may be limited. We performed three additional analyses: 1) IC spectral powers were compared for a contrast between ‘predictive pleasant’ and ‘predictive unpleasant’ to compare against valence and expectation axes; 2) IC spectral powers were compared for a contrast between ‘predictive’ and ‘unpredictive’ to derive a potential predictive axis and compare against the expectation axis; and 3) estimated Waku-Waku was computed and compared against the subjective ratings for each condition.

First, we compared EEGs for a contrast directly contrasting conditions in which Waku-Waku was rated the highest (predictive pleasant) and the lowest (predictive unpleasant). This contrast would not be suited as neural responses for anticipation because this contrast is conceptually subtracting the predictive component out. However, this contrast might provide informative insight on the comparison against the valence (seeing pleasant v.s. seeing unpleasant conditions) or expectation (predictive pleasant v.s. unpredictive) axes. Figure 6 below shows the results of the additional analyses: *θ*-band of IC2, *β*-band of IC7, and *θ* and *α*-band of IC11 were significant (see Supplementary Table for the details).

**Figure 6.**
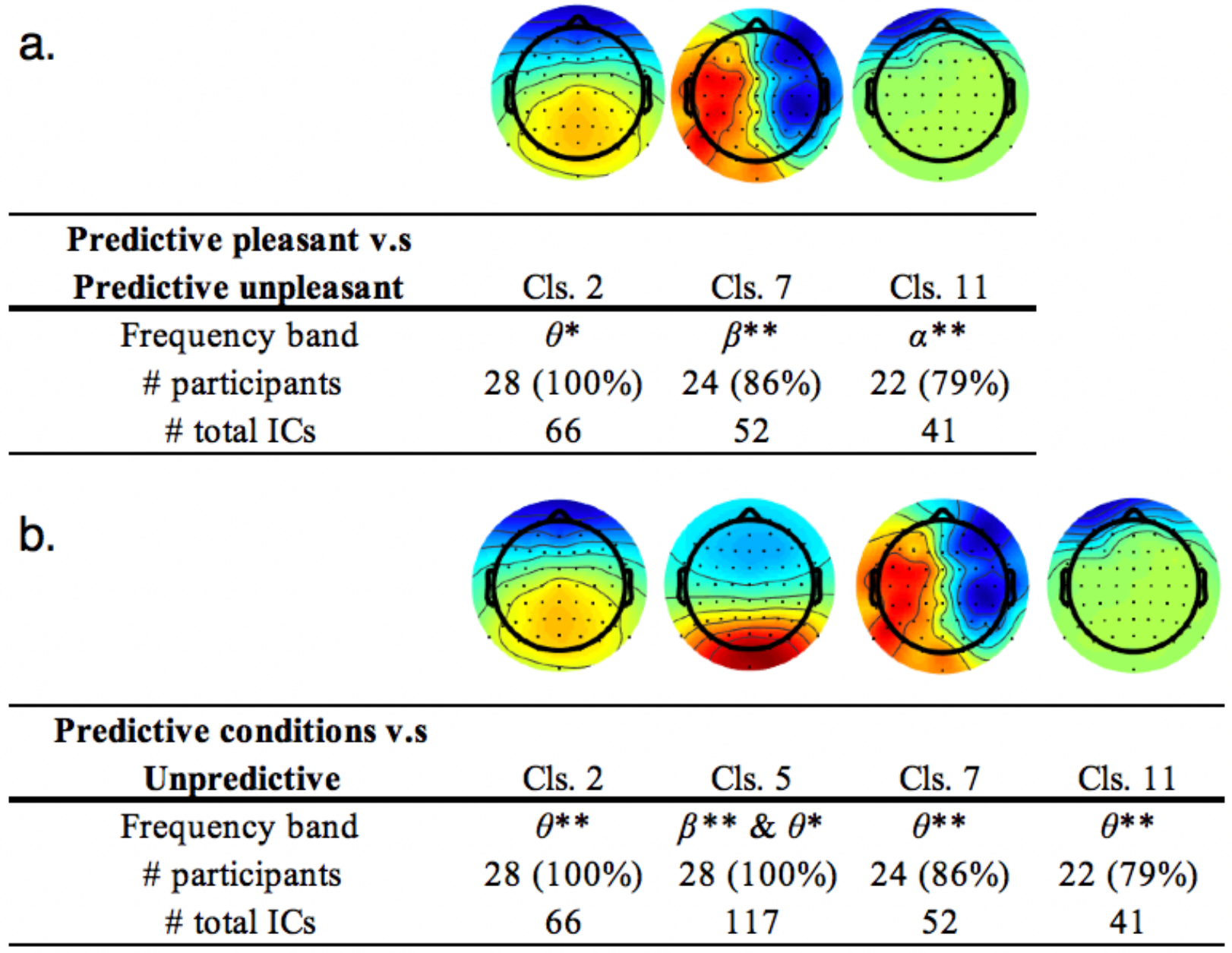
Scalp topographies of significant independent component clusters and their frequency ranges. (a) Conditions in which subjective ratings were the highest and lowest were compared, ‘predictive pleasant’ and ‘predictive unpleasant’, respectively. (b) Conditions of the maximum expectancy regardless of emotional valence (100%) and lowest (50%) expectancy were compared, ‘two predictive conditions combined’ and ‘unpredictive’, respectively. **Statistically significant at *p* < .05 FDR corrected; *Statistically significant at *p* < .05 uncorrected.

To further supplement the inspection, another comparison was also performed on the two predictive conditions combined v.s. unpredictive conditions. In principle, this contrast was supposedly reflecting anticipation of emotionally salient images *regardless* of its emotional value (either pleasant or unpleasant) compared to emotionally vague condition. Therefore, this contrast could have replaced the expectation axis, in theory. As it turned out, *θ*-band of IC2, *β*-band of IC5, *θ*-band of IC6, and *θ*-band of IC11 were found to be significant.

Finally, the proposed BEI model was applied to the existing data by estimating Waku-Waku values based on the given spectra power distributions across the conditions. Based on the assumption, in order to quantify Waku-Waku with the three dimensions, it assumes the existence of the target IC for all axes. However, it turned out that 22 out of 28 participants held all of corresponding ICs in our proposed model; and 24 out of 28 held all corresponding ICs with ‘prediction (IC cluster 5)’ as the third axis.

Table below summarizes the estimated score for each axis and the final Waku-Waku score for each condition. As was expected, using these statistically significant EEG features selected in the formula (4), scores for each axial contrast straightforwardly reflected each pair of contrast (i.e., valence scores for the pleasant image was estimated higher than that for unpleasant, etc.). More importantly, the main candidate of this study, Waku-Waku, seemed well quantified the subjective ratings for each condition, as in Figure 2. Waku-Waku was the highest for the predictive pleasant, followed by unpredictive and predictive unpleasant conditions. Given our proposed model (‘We’) is valid, seeing a highly aroused picture might drive the excitement the most while seeing a picture with low arousal scored the lowest. Please note, only one value per condition is reported, without its mean or standard deviation because this estimation was performed on an average of all EEG features for each axis.

As a counterpart model for the third axis, an alternative model of Waku-Waku was examined by merely replacing the third axis with the ‘prediction’ regardless of emotional valence (a contrast, ‘unpredictable’ v.s. ‘predictive pleasant combined with predictive unpleasant’). As an EEG feature for this axis, *θ*-band of IC cluster 5 was selected as a candidate. The estimated scored did not follow the trend that was achieved by the proposed model. Estimated Waku-Waku scores for the ‘predictive pleasant’ and ‘predictive unpleasant’ was higher than that for the ‘unpredictable’ condition; however, it seemed that the score for the predictive unpleasant was the highest, in which Waku-Waku is supposedly lowest instead.

**Table.** *Estimated* scores for each axis and Waku-Waku for the cue-onset (anticipatory) period and the image-onset period. ‘W_e_’ represents the final estimated Waku-Waku score by the proposed model with an axis of expectation (‘predictive pleasant’ v.s. ‘unpredictive’ conditions) as the third axis. ‘W_p_’ represents Waku-Waku scores estimated by a counterpart model with the third axis replaced by a contrast of prediction (‘two predictive conditions’ v.s. ‘unpredictive’ conditions). The main point of comparison would be the result for the cue-onset, where Waku-Waku and all the subjective ratings were collected. Estimations of Waku-Waku for the image-onset are performed for speculation purposes only. The Waku-Waku scores of the two variant models indicate that our proposed model with the expectation axis outperforms that with the prediction.

|  |  | Valence | Arousal | Expectation | Prediction | We | Wp |
| --- | --- | --- | --- | --- | --- | --- | --- |
| Cue-onset | Predictive pleasant | 54 | 88 | 87 | 41 | 76 | 52 |
|  | Unpredictable | 83 | 26 | 22 | 19 | 46 | 44 |
|  | Predictive unpleasant | 15 | 31 | 36 | 86 | 28 | 54 |
| Image-onset | Pleasant | 89 | 61 | 23 | 86 | 53 | 85 |
|  | Unpleasant | 13 | 41 | 80 | 14 | 51 | 17 |
|  | High arousal | 60 | 88 | 81 | 29 | 75 | 47 |
|  | Low arousal | 35 | 11 | 16 | 72 | 23 | 52 |

## 4. Discussion

We proposed a prototypical model of BEI to quantify “Waku-Waku” towards upcoming visual images using EEG neural markers incorporating a three-dimensional psychological model.

### 4.1 3-D Psychological model

First, a psychological task was given to participants to visually trigger one’s emotion, where participants were required to engage in anticipation of upcoming stimuli. Participants reported their subjective feelings when they were anticipating one of three conditions (predicting pleasant, predicting unpleasant, and unpredictable) on four factors, “Waku-Waku,” valence, arousal, and expectation. As was expected, the 3-D psychological model of “Waku-Waku” with an inclusion of an axis of ‘expectation’ achieved adequately fair fitting accuracy. As quantified by adjusted *R*2 values, the fit of the 3-D model was better by 3 percent than the 2-D model. The improvement of the fit may be trivial to our aim, but at least the multi-axial model of emotion may be feasible to adapt.

Also, the 3-D model revealed a high loading on the third axis relative to the rest of the two axes that the classical 2-D circumplex model of emotion would propose [1]. It assures that the subjective feeling of “Waku-Waku” may intimately link to anticipation, and followed by momentary emotions of pleasure and arousal. We have also found the two subjective ratings of valence and expectation moderately correlated with that of Waku-Waku, yet they were not equal (the power of correlation (*r*2) was not very strong). It also suggests that Waku-Waku and expectation-only or valence-only were not the same, but somewhat reflecting a subset of Waku-Waku feeling. This result was plausible because its definition of “Waku-Waku” is described as a state of one’s heart is moved due to being pleased and expecting something pleasant. Therefore, both valence and expectation may be assumed to be highly loaded. Nevertheless, as was discussed earlier, *Kansei*, or our instantaneous mental state of emotion may be modeled by multifold of human affects and cognition [6-8,11]. Furthermore, the resultant variations in coefficients for valence, arousal, and expectation would suggest that the three aspects of psychological dimensions differentially contribute to the feeling of Waku-Waku.

A psychological model shall not be restricted only by these proposed three axes (valence, arousal, and expectation). Particularly the concept of the third axis in our case was a concept of time-domain; however, any other dimensions associated with human senses may be acquired. As represented by the PAD (pleasure, activation, and dominance) model, dominance [37] might have been the third axis. As discussed later, a sense of prediction that may putatively reflect a likelihood of expectation may also be a key factor because it is rooted in the concept of predictive coding. Furthermore, it has been argued that a psychological constructionist approach assumes emotion as the interplay of the multiple brain networks that may require potentially more than three dimensions or other sophisticated methods [6]. Nevertheless, it is plausible to propose that modeling of our putatively complex nature of awareness would benefit from the multi-dimensional model.

### 4.2 Corresponding electrophysiological markers for the three axes

As to determine neural correlates of each axis of BEI, spectral powers of ICs of EEGs were analyzed. Valence and arousal axes were quantified from the duration in which participants visually saw an emotionally triggering picture. Expectation axis was determined from the delay period in which they were anticipating upcoming picture according to played auditory cue tied to valence types. A conventional spectral power analysis was performed on each IC. Dissociable neural markers were identified for each axis. This dissociation is one of the essential aspects for the validation of our model. Because the multi-dimensional model assumes independence of psychological axes and their corresponding neural markers, it would be necessary to confirm EEG features selected for each axis would be dissociable. For example, a previous study suggests that multiple ICs would be loaded on one axis and that one IC might also contributing to the other axes to some extent [39]. In addition, studies support a view that the functions of a brain region are ubiquitous and not limited to a unitary and discrete function [6]. Therefore, we do not necessarily stringent on this overlap; however, a complete overlap between a pair would be another story. If the same EEG features entirely correspond to a pair of axes, it may suggest the pair would be electrophysiologically an identical system. As a result, it might violate the assumption of the multi-dimensional model, making the addition of a new dimension meaningless. However, our results declined this possibility. There was no complete overlap across the three axes, supporting our proof of concept. The three-dimensional model, particularly the newly added dimension, would plausibly reflect putatively dissociable neural mechanisms. Below, we discuss the outcomes for each axis.

#### 4.2.1 IC markers of valence

For the valence axis, among several significant EEG features (*θ*-band of IC1, *θ* & *α*-bands of IC5, α-band of IC6, *θ* & β-bands of IC7and β-band of IC14), *θ*-band of IC cluster 5 was robustly significant, and the component was found in all (100%) of participants. Previous neuroimaging research suggests a source in proximity to orbitofrontal areas may be responsible for emotional valence [38]. A similar EEG study [39] related EEG features to the subjective feeling at rest, rather than during a task based on another type of 3-D emotional space (valence, arousal, and dominance, also known as the PAD model). By focusing only on the *<u>β</u>*<u>-band</u> power of IC clusters, they found that IC clusters with sources localized in posterior cingulate and right posterior temporal lobe were positively correlated with valence. In our study, an estimated dipole of clusters 6 and 7 centered around the mid-to posterior cingulate regions in close proximity to their finding (see Supplementary Figure 2). Notably, *β*-band of cluster 7 was also significant, validating the replicability of this neural source for emotional valence. However, in our case, because the number of participants who held this component was relatively small, we selected the alternative target with its significant frequency band in *<u>θ</u>*<u>-band</u> instead. This IC could have been a reliable candidate for valence if the sampled population differ. Our finding may indicate relatively slow oscillation at *θ*-band that might be induced by processing visual information, and such slow oscillatory activity may be associated with remotely interconnected reward networks such as midline orbitofrontal and anterior cingulate regions reported in fMRI studies [13].

#### 4.2.2 IC markers for arousal

As for the arousal axis, neural activities evoked by seeing a picture with *high* arousal (i.e., a picture of fireworks or explosion, etc.) were compared against that with *low* arousal (i.e., a picture of a calm scenery of house or a kitten, etc.). Just because of a slight difference in reliability across participants, *α*-band of IC cluster 7 was selected as the best target. Whereas another prominent candidate was *β*-band of IC11, this component might have been selected as an alternative. As in convention, *α*-band oscillations elicited from parietal regions typically reflect human arousal [22-24]. Our results would suggest that *α*-band activities with potential sources around the midline part of the brain (cingulate cortex) may reflect arousal. While we carefully selected pictures based on their visuophysical properties, yet some physical features of pictures such as luminance or brightness instead of their content might covertly trigger some of EEG signals unrelated to the subjective feeling of arousal. Future studies may need to investigate on this to rule out some potentially confounding factors.

#### 4.2.3 IC markers for expectation

As for the expectation axis, comparisons of neural activities when expecting a pleasant picture against when valence of expecting picture was unknown, *θ*-band of IC cluster 10 with its dipole centered around right angular gyrus (proximity to inferior parietal lobule and lateral occipital complex regions) was significant. This region is known to be responsible for visuospatial attention [40] or maintenance of visual information in memory [41]. Another candidate was *α*-band cluster 15 with its localized source around the dorsolateral prefrontal cortex (DLPFC), typically nominated as a central source of executive functions [42,43]; however, accumulating research also suggest that this region would be playing a pivotal role in the integration of emotion and cognition [13]. Notably, studies of depression and emotional valence sometimes refer to the *α*-band hemispheric asymmetry with its sources in this prefrontal region [44,45], and this EEG feature is widely applied in BCI research aiming for treatment of depression [46,47].

Again, it could have been possible that we might observe the same or similar neural marker for expectation as that for valence axis because they merely differed whether they were anticipating in mind or seeing a picture in front of them. One may speculate that contrasts for the valence and the expectation might overlap. However, the similar approach of showing emotional pictures while brain functions are monitored in fMRI [48], Bermpohl, et al. (2006) reported lateral occipital regions are activated while *seeing* emotional pictures rather than expecting phase, while anterior and posterior cingulate regions were responsible for expectation compared to neutral targets. One may argue that methodological differences between EEG and fMRI, as EEGs may be suitable for detecting electrical discharges while fMRI tracks cerebral blood flows. Another putative explanation may be that in our comparison, we did not have pictures with neutral valence. In our design, even at the unpredictable condition, anticipated imagery and subjective rating for this condition were fluctuating between the two extremities of pleasant and unpleasant across participants (see Supplementary Figure2); therefore, this unpredictable condition would not be same as expecting a neutral image. Detailed investigations would be necessary to discuss further about the overlap between the location of dipoles with BOLD responses obtained from fMRI studies.

Nonetheless, previous fMRI research supports dissociable networks for emotional expectancy and emotion perception [48], corresponding to the expectation and the valence axes, respectively. As for EEG studies on expectancy, several neural markers such as late positive potentials, readiness potentials, or some other preparatory EEG markers reportedly influence the expectancy of emotional items [49-51]. In our results, expectancy processes might recruit a combination of the executive network and visuospatial network to anticipate by forming imagery in mind. Our findings on this axis may contribute to the field of expectancy. In reality, a sophisticated interplay of these higher cognitive functions may be associated with the expectation of emotional events. Notably, the selected neural marker for the expectation axis was also dissociable against the ones found to be significant for the valence axis. This neural level of dissociation indirectly assures that these two psychological dimensions would also be distinct, even the psychological ratings of the two correlated to some extent. It may support that the inclusion of the expectation axis may benefit from quantifying dissociable neural processes underlying the ubiquitous interplay of cognitive or affective functions [6].

### 4.3 Alternative comparisons and validation of the BEI model

In this study, subjective ratings of Waku-Waku and the other axial factors were not collected at every trial. However, as an attempt to validate the neural markers found in the three axes, we directly contrasted the conditions in which Waku-Waku ratings were the highest (‘predicting pleasant’) and lowest (‘predicting unpleasant’). Conceptually speaking, the valence axis, and this contrast might have shared the same EEG features because both contrasts are putatively comparing emotionally pleasant and unpleasant images. The only difference between the two was whether participants were seeing an actual image on the screen or forming a mental image in mind. However, no EEG features overlapped between the two, suggesting the neural mechanisms associated with the emotional valence of expecting a future and that of exogenously confronting one may be dissociable. To iterate, in order to estimate Waku-Waku, the inclusion of anticipation was a fundamental aspect of our model. Therefore, it was presumed that EEG features selected here would not stand the expectation axis because it would eliminate this valuable perspective of anticipatory neural function by contrasting the two predictive conditions.

Another potential contrast was made on ‘two predictive conditions combined v.s. unpredictive’. Several neural signatures emerged on this axis, and they could become a counterpart for the third axis of expectation. The selected EEG feature for predictability successfully quantified the anticipatory conditions to some extent. However, the estimation of Waku-Waku (W_p_) with this axis failed to follow the subjective ratings. A close look on the Table indicates that the predictive axis rated the highest (‘86’) when the cue was predictive *unpleasant* whereas the estimated prediction value for the predictive pleasant, it was only (‘41’). It might have been true that participants might “predicted” an occurrence of unwanted images; however, this contrast was not satisfactory to represent the ‘expectation’ axis. Albeit the fit was not great for Waku-Waku, it does not necessarily mean that this axis of prediction shall not be a part of the multi-axial model of emotion. It may be possible that the neural responses for predictive unpleasant (somewhat associated with an emotion of worry) might be stronger than that for predictive pleasant. If the experimental design and the way we asked the participants to report was more closely related to the prevalence, or in another word, “entropy” of expectation regardless of emotional value, such axis and the selected components might have been acquired as a right candidate as prediction or certainty axis. Altogether, this axis was not suitable for our proposed model for Waku-Waku. Finally, it would be plausible to believe that the EEG feature we selected for the contrast between the ‘predictive pleasant’ and ‘unpredictive’ was appropriate for our model.

## 5. Conclusion

We proposed a prototype of BEI based on a multi-axis, 3-dimensional model of emotion to quantify our anticipatory excitement using EEG. Fidelity of the BEI shall be examined in future studies; however, provided a certain degree of accuracy backed by statistical results, our BEI may be able to quantify and applicable at least for young adult Japanese (or Asian) individuals. In our group-level analysis, we found only one IC and its corresponding frequency band for valence and expectation axes; however, our result may not be conclusive due to putative cultural or age differences. Also, as our EEG-based BEI model proposed only for visual stimuli, similar experiments or its generalization need to be tested with stimuli on other modalities, such as audition and tactile.

Moreover, we found that the number of ICs shared by our participants were not perfect, especially not all participants shared some of the key IC clusters selected for arousal and expectation axes. It implies that the currently proposed BEI model may not be generalized for all individuals, even within our collected samples. A close investigation and individual optimization for selecting IC and its frequency range may be necessary to achieve full compatibility of the BEI. We applied the GMM method to determine the number of ICs to extract. While this method may be a quantitative means to determine the number of clusters, this approach tends to fluctuate using slightly different parameters. Alternatively, it may simplistically differ in a different ethical, cultural, or age population. Therefore, one should be careful when applying a result observed here. Based upon a fixed-effect model by determining a group-average model, we selected a fixed EEG feature across participants. Recent studies of neuroscience instead propose individually optimized decoding of neural activities outperform a group-level approach [52,53], potentially associated with neuroarchitectures that differ across age [54,55] or personality [56-58]. It is indeed plausible that individual-wise optimization of either or both psychological and neural models may provide accurate quantification of our feelings.

Our observations in this article may be limited in various aspects; however, this should constitute a reasonable basis to quantify our sense of *Kansei*. There are wide varieties of BCIs that exist in the field, our approach of considering multiple axes combined with EEG markers may become a new tool for a neuroscientific consultation. Such a tool may be applicable not only for stable pictures (i.e., seeing an art, picture, advertisement posters, etc) but also be useful for various other situations, such as evaluating emotional responses for seeing a motion-pictures (movies, TV commercials). BEI may certainly require further evidence and theoretical supports; however, it may become a useful tool for *Kansei* engineering in the near future.

## Acknowledgements

Patents (PCT/JP2016/003712; JP2016116449A; US15/768,782; EP16855084.6A) have been submitted based on a part of described methods in this manuscript.

All authors designed behavioral paradigm and discussed the manuscript. N.K. and K.M. prepared experimental materials and collected data. MGM, N.K., and R.M. performed the EEG analyses. M.G.M. and G.L. designed and built the brain computer interface. MGM, NK, and GL wrote the manuscript. We highly appreciate Prof. Hirokazu Yanagihara at Hiroshima University for his mathematical advice on this project.

This research was supported by JST COI Grant Number JPMJCE1311. GL was supported by the New Energy and Industrial Technology Development Organization (NEDO), by ImPACT of CSTI and by the Commissioned Research of NICT.

