## Supplementary Materials for "Quantification of anticipation of excitement with three-axial model of emotion with EEG"

### Supplementary Table 1

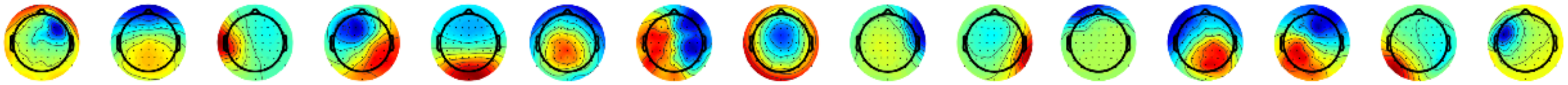

| | Cls. 1 | Cls. 2 | Cls. 3 | Cls. 4 | Cls. 5 | Cls. 6 | Cls. 7 | Cls. 8 | Cls. 9 | Cls. 10 | Cls. 11 | Cls. 12 | Cls. 13 | Cls. 14 | Cls. 15 |
|---|---|---|---|---|---|---|---|---|---|---|---|---|---|---|---|
| # participants | 26 (93%) | 28 (100%) | 24 (86%) | 25 (89%) | 28 (100%) | 26 (93%) | 24 (86%) | 23 (82%) | 25 (89%) | 25 (89%) | 22 (79%) | 28 (100%) | 25 (89%) | 26 (93%) | 24 (86%) |
| # total ICs | 43 | 66 | 59 | 75 | 117 | 62 | 52 | 79 | 51 | 44 | 41 | 66 | 78 | 66 | 50 |
| **Valence (pleasant v.s. unpleasant)** | | | | | | | | | | | | | | | |
| θ | **2.25 ( .07 )** | -1.29 ( .59 ) | -0.62 ( .53 ) | -0.90 ( .54 ) | **2.39 ( .03 )** | 1.13 ( .25 ) | **2.28 ( .05 )** | -0.06 ( .95 ) | -0.84 ( .39 ) | 0.65 ( .77 ) | 0.90 ( .67 ) | 1.11 ( .40 ) | 0.19 ( .85 ) | 1.49 ( .13 ) | 0.99 ( .74 ) |
| α | -0.08 ( .93 ) | 0.16 ( .87 ) | 0.63 ( .53 ) | -1.01 ( .54 ) | **2.23** ( **.03** ) | **2.81** ( **.01** ) | 0.32 ( .74 ) | 0.89 ( .56 ) | 0.90 ( .39 ) | 0.81 ( .77 ) | -0.42 ( .67 ) | 1.57 ( .35 ) | 0.41 ( .85 ) | 1.87 ( .09 ) | 0.38 ( .74 ) |
| β | 1.04 ( .44 ) | -0.40 ( .87 ) | 1.35 ( .53 ) | -0.22 ( .82 ) | 1.55 ( .12 ) | 1.27 ( .25 ) | **2.11 ( .05 )** | 1.47 ( .42 ) | -0.96 ( .39 ) | 0.04 ( .97 ) | -0.72 ( .67 ) | 0.32 ( .75 ) | 2.04 ( .12 ) | **2.53** ( **.03** ) | -0.32 ( .74 ) |
| **Arousal (high v.s. low)** | | | | | | | | | | | | | | | |
| θ | -0.76 ( .83 ) | 0.96 ( .41 ) | -0.20 ( .84 ) | 0.37 ( .70 ) | 0.40 ( .99 ) | -1.54 ( .37 ) | -0.52 ( .90 ) | 0.35 ( .72 ) | -0.57 ( .56 ) | -0.63 ( .52 ) | 0.38 ( .70 ) | -1.78 ( .19 ) | -1.32 ( .56 ) | 0.00 ( .99 ) | 0.15 ( .88 ) |
| α | -0.41 ( .83 ) | 1.16 ( .41 ) | 1.17 ( .55 ) | 0.82 ( .61 ) | 0.90 ( .99 ) | -0.54 ( .59 ) | **2.01 ( .13 )** | 0.58 ( .72 ) | 1.84 ( .19 ) | -0.82 ( .52 ) | 1.04 ( .44 ) | -1.52 ( .19 ) | -0.52 ( .60 ) | 1.87 ( .18 ) | 0.84 ( .88 ) |
| β | -0.21 ( .83 ) | 0.81 ( .41 ) | -0.89 ( .55 ) | 1.47 ( .42 ) | 0.00 ( .99 ) | 0.54 ( .59 ) | -0.05 ( .95 ) | 1.95 ( .15 ) | 1.30 ( .28 ) | -0.95 ( .52 ) | **2.44 ( .04 )** | -0.31 ( .75 ) | 0.73 ( .60 ) | -0.75 ( .67 ) | -0.48 ( .88 ) |
| **Expectation (predicting pleasant v.s. predicting unpleasant)** | | | | | | | | | | | | | | | |
| θ | 1.00 ( .63 ) | -1.31 ( .40 ) | -1.86 ( .18 ) | -0.57 ( .57 ) | -0.53 ( .59 ) | -1.24 ( .52 ) | -0.29 ( .77 ) | -0.30 ( .90 ) | -1.21 ( .67 ) | **-2.05 ( .11 )** | -0.55 ( .59 ) | -0.14 ( .89 ) | -0.23 ( .85 ) | 0.51 ( .63 ) | -0.09 ( .95 ) |
| α | 0.47 ( .63 ) | -1.11 ( .40 ) | -0.94 ( .44 ) | 0.62 ( .57 ) | -0.74 ( .59 ) | -0.42 ( .67 ) | 0.85 ( .77 ) | 0.12 ( .90 ) | 0.35 ( .72 ) | -0.72 ( .70 ) | 0.52 ( .59 ) | -0.41 ( .89 ) | -0.18 ( .85 ) | 0.69 ( .63 ) | **2.18 ( .08 )** |
| β | 0.80 ( .63 ) | 0.59 ( .55 ) | -0.77 ( .44 ) | -0.73 ( .57 ) | -1.87 ( .18 ) | -0.94 ( .52 ) | 0.40 ( .77 ) | 0.52 ( .90 ) | 0.60 ( .72 ) | 0.22 ( .82 ) | 1.08 ( .59 ) | -1.94 ( .15 ) | -0.73 ( .85 ) | 0.47 ( .63 ) | -0.05 ( .95 ) |
| **Predicting pleasant v.s. predicting unpleasant** | | | | | | | | | | | | | | | |
| θ | 1.91 ( .09 ) | **3.38 ( .00 )** | -1.19 ( .35 ) | 0.19 ( .91 ) | 0.99 ( .33 ) | 1.66 ( .29 ) | 1.16 ( .37 ) | 1.53 ( .18 ) | -0.14 ( .88 ) | -0.83 ( .61 ) | **2.82 ( .01 )** | 1.45 ( .22 ) | 0.92 ( .75 ) | -0.90 ( .74 ) | 0.17 ( .86 ) |
| α | 1.23 ( .21 ) | 0.98 ( .43 ) | -1.55 ( .35 ) | 0.11 ( .91 ) | -0.96 ( .33 ) | -0.77 ( .52 ) | 0.53 ( .59 ) | 0.54 ( .59 ) | -1.34 ( .27 ) | -0.08 ( .93 ) | **2.69 ( .01 )** | 1.03 ( .30 ) | 0.31 ( .75 ) | -0.32 ( .74 ) | 1.30 ( .58 ) |
| β | 1.86 ( .09 ) | 0.78 ( .43 ) | -0.02 ( .98 ) | 0.19 ( .91 ) | 1.50 ( .33 ) | 0.63 ( .52 ) | **2.43 ( .04 )** | 1.62 ( .18 ) | -1.40 ( .27 ) | 1.74 ( .24 ) | -0.60 ( .54 ) | -1.82 ( .20 ) | 0.31 ( .75 ) | 0.39 ( .74 ) | -0.69 ( .73 ) |
| **Unpredictable v.s. Predictable (predicting pleasant and predicting unpleasant)** | | | | | | | | | | | | | | | |
| θ | 1.15 ( .27 ) | **4.09 ( .00 )** | 0.83 ( .64 ) | 0.88 ( .56 ) | **1.54 ( .18 )** | **1.94 ( .10 )** | **2.42 ( .04 )** | 2.03 ( .12 ) | 1.23 ( .26 ) | 0.50 ( .61 ) | **3.66 ( .00 )** | 1.66 ( .21 ) | 1.71 ( .23 ) | -1.64 ( .29 ) | 0.34 ( .73 ) |
| α | 1.12 ( .27 ) | 1.45 ( .21 ) | -0.13 ( .89 ) | -0.25 ( .79 ) | -0.31 ( .75 ) | -0.89 ( .37 ) | -0.22 ( .82 ) | 0.25 ( .80 ) | -1.12 ( .26 ) | 0.93 ( .61 ) | 1.13 ( .25 ) | 1.24 ( .21 ) | 0.27 ( .78 ) | -0.56 ( .57 ) | -0.72 ( .70 ) |
| β | 1.09 ( .27 ) | -0.05 ( .96 ) | 0.79 ( .64 ) | 1.18 ( .56 ) | **3.18 ( .00 )** | 1.81 ( .10 ) | 1.64 ( .15 ) | 1.43 ( .23 ) | -1.27 ( .26 ) | 0.55 ( .61 ) | -1.65 ( .14 ) | 1.47 ( .21 ) | 1.42 ( .23 ) | 0.74 ( .57 ) | -1.23 ( .65 ) |

Summary of clustered ICs. Number of participants and percentage of participants among all who held an IC clustered into each IC cluster (“Cls. 1” to “Cls. 15”). All statistical results (*Z*-scores and FDR corrected *p*-values in parentheses) of Wilcoxon signed-rank test on each frequency range are shown for all comparisons of interest. **Red bold** texts represent robustly significant frequency at $p_{\text{FDR-corrected}} < .05$; **bold texts** represent significant frequency at $p_{\text{uncorrected}} < .05$; and underlined texts represent marginally significant frequency at $p_{\text{uncorrected}} < .10$. As was obvious, different IC and frequency ranges reflected different aspects of the psychological axis, valence, arousal, and expectation. In addition, three IC clusters out of fifteen appeared to be shared by all (100%) participants, suggesting such IC may be applicable for anyone; however, some IC clusters was not shared by all (i.e. IC cluster 11 for valence and IC cluster 10 were shared by only 79% and 89%, respectively). As a supplement to the main axes of interest (valence, arousal, and expectation), two additional comparisons are presented for the sake of completeness: a contrast between predicting pleasant v.s. predicting unpleasant and contrast between predictive conditions (concatenating both predictive pleasant and predicting unpleasant) v.s. unpredictable condition

### Supplementary Figure 1

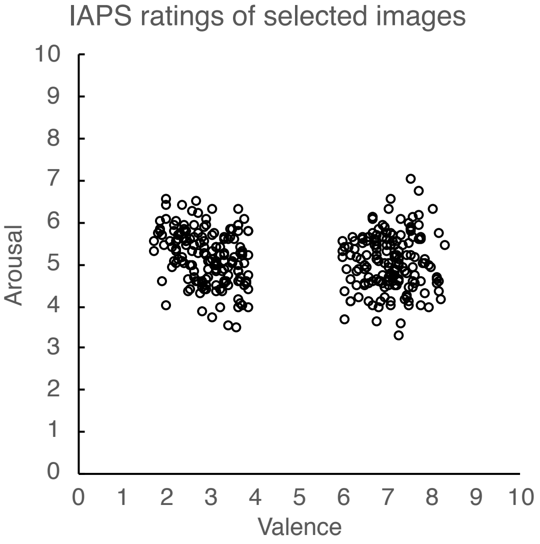


A scatter plot of subjective ratings (as reported in the Lang, Bradley, & Cuthbert, 2008) selected in this study for valence (along the X-axis) and arousal (along the Y-axis). Pictures with intermediate valence (with valence rating between 4–6) were not used in this study. As is apparent in the figure below, we selected pictures so that the valence cue would be congruent to its assignment: one for pleasant (valence value above 6) and the other for unpleasant (valence value below 4). As for *arousal*, we did not set such a border, yet simply applied a median split for high and low arousal pictures. See Supplementary Table 2 below for the details (mean ± *SD*) of valence and arousal ratings for selected sets of pictures.

### Supplementary Table 2

| | | Set 1 | Set 2 |
|---|---|---|---|
| Valence | Pleasant | 7.17 ± 0.53 | 6.99 ± 0.57 |
| | Unpleasant | 3.06 ± 0.61 | 2.91 ± 0.55 |
| Arousal | High | 5.31 ± 0.45 | 5.77 ± 0.28 |
| | Low | 4.41 ± 0.27 | 4.86 ± 0.52 |

### Supplementary Figure 2

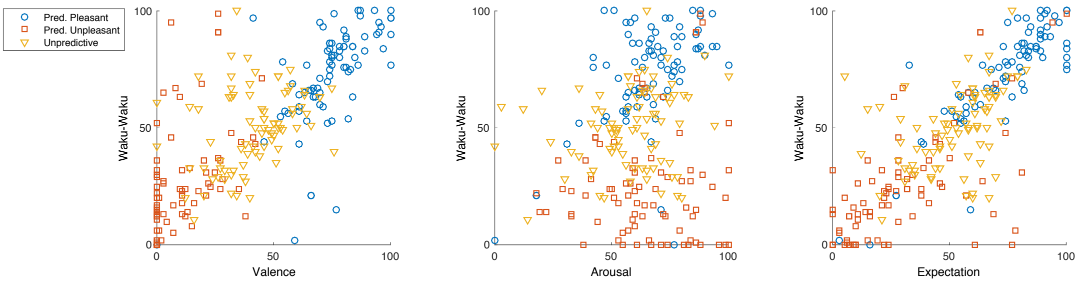


The figure shows scatter plots of Waku-Waku values as a function of valence (left panel), arousal (middle panel) and expectation (right panel). Each of 3 conditions is plotted separately: blue circles for predictive pleasant, red squares for predictive unpleasant, and yellow triangles for unpredictive conditions. These correlations for valence and expectation were significant; however, the power of the results was moderately high for valence and expectation axes, and not as very strong nor perfect. The direct correlation was weak to negligible for arousal, yet it was significant by the inclusion of 9 samples per participant (3 conditions each for 3 sessions). While these are correlated, derived coefficients by the mixed model take these correlations and innate subject-effects into account.

### Supplementary Figure 3

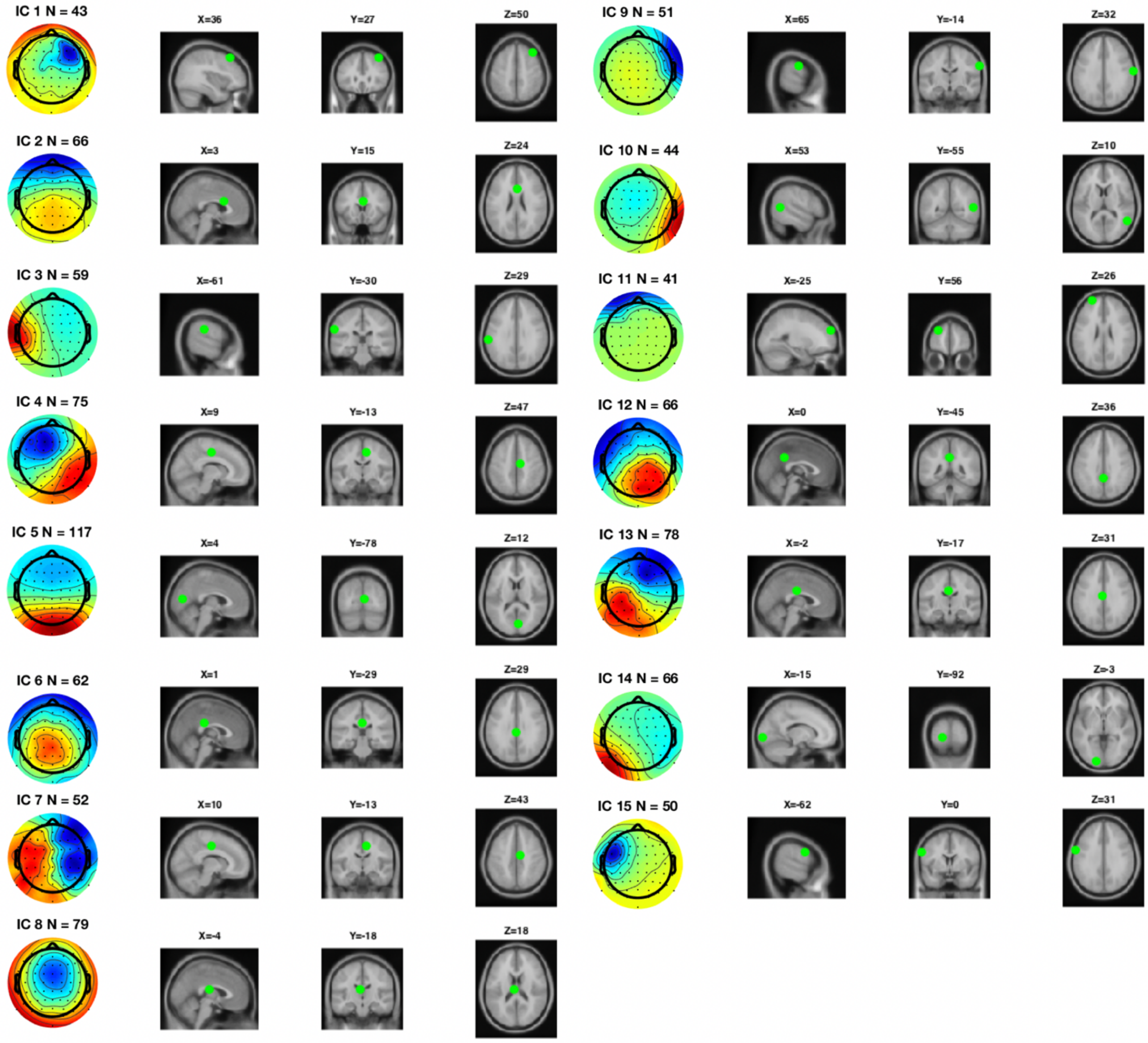


Scalp topography of each IC cluster with the number of ICs in the cluster and a location of the centroid (green dot) plotted on a template brain for each IC cluster with its spherical coordinates. Centroids for IC5, 7, and 10 (selected for valence, arousal, and expectation) were centered around posterior cingulate cortex to precuneus, visual cortex in proximity to posterior cingulate and precuneus regions, and right angular gyrus (lateral occipital complex) regions, respectively. For each IC, potential brain regions were identified according to their MNI coordinates of centroid based on a standardized AAI brain template, as follows. If a centroid is at a border of two regions, both are listed. IC1: right mid-frontal cortex; IC2: right anterior cingulate cortex; IC3: left supramarginal gyrus; IC4: right mid-cingulate cortex; IC5: precuneus, calcarine (primary visual cortex); IC6: right mid-cingulate cortex; IC7: right mid-cingulate cortex; IC8: left thalamus; IC9: right postcentral gyrus; IC10: right angular gyrus, mid-temporal gyrus; IC11: left mid-frontal cortex; IC12: mid-cingulate cortex, precuneus; IC13: left mid-cingulate cortex; IC14: left superior occipital cortex; IC15: left prefrontal gyrus, postcentral gyrus.

**Appendix**: 320 IAPS picture codes acquired from the IAPS database

| | | | | | | | |
|---|---|---|---|---|---|---|---|
| 1090 | 2075 | 2387 | 4007 | 5628 | 7481 | 9001 | 9390 |
| 1111 | 2091 | 2389 | 4090 | 5820 | 7482 | 9002 | 9403 |
| 1113 | 2095 | 2395 | 4150 | 5829 | 7489 | 9008 | 9404 |
| 1220 | 2100 | 2398 | 4225 | 5830 | 7492 | 9031 | 9409 |
| 1270 | 2110 | 2399 | 4250 | 5831 | 7499 | 9041 | 9417 |
| 1270 | 2115 | 2455 | 4255 | 5833 | 7502 | 9043 | 9421 |
| 1271 | 2120 | 2456 | 4532 | 5910 | 7521 | 9046 | 9425 |
| 1274 | 2141 | 2457 | 4597 | 5961 | 7530 | 9050 | 9426 |
| 1275 | 2150 | 2530 | 4598 | 5973 | 7660 | 9090 | 9427 |
| 1280 | 2152 | 2590 | 4599 | 5994 | 8001 | 9102 | 9428 |
| 1301 | 2153 | 2655 | 4600 | 6240 | 8021 | 9110 | 9429 |
| 1340 | 2155 | 2661 | 4601 | 6242 | 8031 | 9140 | 9430 |
| 1410 | 2158 | 2683 | 4603 | 6244 | 8034 | 9145 | 9435 |
| 1440 | 2160 | 2694 | 4610 | 6311 | 8090 | 9180 | 9440 |
| 1441 | 2165 | 2700 | 4612 | 6530 | 8130 | 9182 | 9445 |
| 1460 | 2170 | 2703 | 4614 | 6561 | 8161 | 9184 | 9470 |
| 1463 | 2205 | 2715 | 4617 | 6831 | 8162 | 9185 | 9471 |
| 1540 | 2208 | 2716 | 4619 | 6838 | 8163 | 9186 | 9490 |
| 1590 | 2209 | 2722 | 4621 | 7079 | 8190 | 9190 | 9491 |
| 1601 | 2216 | 2745.2 | 4622 | 7135 | 8193 | 9220 | 9495 |
| 1620 | 2217 | 2750 | 4623 | 7200 | 8208 | 9230 | 9530 |
| 1630 | 2222 | 2753 | 4624 | 7240 | 8210 | 9265 | 9560 |
| 1659 | 2224 | 2795 | 4626 | 7250 | 8230 | 9270 | 9561 |
| 1710 | 2274 | 2799 | 4628 | 7260 | 8231 | 9280 | 9571 |
| 1720 | 2276 | 2800 | 4640 | 7279 | 8330 | 9290 | 9592 |
| 1721 | 2278 | 2900.1 | 4641 | 7280 | 8340 | 9291 | 9594 |
| 1750 | 2300 | 2900.2 | 4645 | 7284 | 8370 | 9295 | 9610 |
| 1810 | 2301 | 3022 | 4700 | 7286 | 8380 | 9300 | 9622 |
| 1812 | 2303 | 3180 | 5199 | 7330 | 8420 | 9301 | 9630 |
| 1920 | 2306 | 3181 | 5215 | 7350 | 8461 | 9302 | 9800 |
| 2019 | 2311 | 3185 | 5250 | 7352 | 8467 | 9320 | 9831 |
| 2030 | 2312 | 3195 | 5270 | 7359 | 8470 | 9322 | 9900 |
| 2039 | 2314 | 3215 | 5450 | 7360 | 8480 | 9325 | 9903 |
| 2040 | 2340 | 3216 | 5470 | 7361 | 8485 | 9326 | 9912 |
| 2050 | 2345 | 3220 | 5480 | 7380 | 8496 | 9330 | 9920 |
| 2055.1 | 2345.1 | 3230 | 5551 | 7390 | 8499 | 9331 | 9921 |
| 2057 | 2346 | 3280 | 5621 | 7430 | 8500 | 9332 | 9922 |
| 2058 | 2347 | 3300 | 5622 | 7470 | 8510 | 9341 | 9925 |
| 2070 | 2352.1 | 3350 | 5623 | 7477 | 8531 | 9342 | 9930 |
| 2071 | 2375.1 | 4006 | 5626 | 7480 | 8540 | 9373 | 9941 |
